# Species-level bacterial community profiling of the healthy sinonasal microbiome using Pacific Biosciences sequencing of full-length 16S rRNA genes

**DOI:** 10.1101/338731

**Authors:** Joshua P. Earl, Nithin D. Adappa, Jaroslaw Krol, Archana S. Bhat, Sergey Balashov, Rachel L. Ehrlich, James N. Palmer, Alan D. Workman, Mariel Blasetti, Bhaswati Sen, Jocelyn Hammond, Noam A. Cohen, Garth D. Ehrlich, Joshua Chang Mell

## Abstract

**Background:** Pan-bacterial 16S rRNA microbiome surveys performed with massively parallel DNA sequencing technologies have transformed community microbiological studies. Current 16S profiling methods, however, fail to provide sufficient taxonomic resolution and accuracy to adequately perform species-level associative studies for specific conditions. This is due to the amplification and sequencing of only short 16S rRNA gene regions, typically providing for only family- or genus-level taxonomy. Moreover, sequencing errors often inflate the number of taxa present. Pacific Biosciences’ (PacBio’s) long-read technology in particular suffers from high error rates per base. Herein we present a microbiome analysis pipeline that takes advantage of PacBio circular consensus sequencing (CCS) technology to sequence and error correct full-length bacterial 16S rRNA genes, which provides high-fidelity species-level microbiome data

**Results:** Analysis of a mock community with 20 bacterial species demonstrated 100% specificity and sensitivity. Examination of a 250-plus species mock community demonstrated correct species-level classification of >90% of taxa and relative abundances were accurately captured. The majority of the remaining taxa were demonstrated to be multiply, incorrectly, or incompletely classified. Using this methodology, we examined the microgeographic variation present among the microbiomes of six sinonasal sites, by both swab and biopsy, from the anterior nasal cavity to the sphenoid sinus from 12 subjects undergoing trans-sphenoidal hypophysectomy. We found greater variation among subjects than among sites within a subject, although significant within-individual differences were also observed. *Propiniobacterium acnes* (recently renamed *Cutibacterium acnes* [1]) was the predominant species throughout, but was found at distinct relative abundances by site.

**Conclusions:** Our microbial composition analysis pipeline for single-molecule real-time 16S rRNA gene sequencing (MCSMRT, https://github.com/jpearl01/mcsmrt) overcomes deficits of standard marker gene based microbiome analyses by using CCS of entire 16S rRNA genes to provide increased taxonomic and phylogenetic resolution. Extensions of this approach to other marker genes could help refine taxonomic assignments of microbial species and improve reference databases, as well as strengthen the specificity of associations between microbial communities and dysbiotic states.

## Background

The advent of culture- and cloning-free methods to analyze bacterial phylogenetic marker genes by deep sequencing ushered in a new era of microbial community analysis, dramatically reducing the labor and cost of profiling the identities and abundances of microbes from different environments, independent of their ability to be cultivated [2-5]. The small subunit ribosomal RNA gene (16S rRNA) is shared by all bacteria and has been sequenced in thousands of distinct named species. Because of this, polymerase chain reactions (PCR) using primers that target conserved regions can amplify variable segments of the 16S rRNA gene from across the bacterial domain in a relatively unbiased fashion for amplicon-based deep sequencing [6, 7]. 16S sequence databases can then be used to classify a given sequence read’s taxonomic source. Combined with increasingly powerful ecological methods for analyzing microbial community dynamics and inferring community-level metabolic networks, profiling the taxonomic composition of bacterial communities by 16S rRNA gene sequencing has become a standard part of microbiome analysis [8-11].

Unfortunately, the use of the 16S rRNA gene as a taxonomic marker has, in part, been constrained by the short read length of the most commonly used sequencing platform for microbial community profiling (the Illumina MiSeq), which only allows interrogation of up to 3 of 9 variable regions in the 16S rRNA gene (called V1-V9), often targeting only V3-V5, V1-V3, or V4 alone [10, 12-16]. This constraint limits the taxonomic resolution to which short reads can be classified, typically only to the family- or genus-level, and furthermore taxonomic resolution varies for different groups of bacteria when using different portions of the 16S rRNA gene [10]. Low-resolution classification in turn limits not only the accuracy and precision of ecological inferences and metabolic reconstructions, but also the ability to identify appropriate bacterial strains to use in follow-up experimental and translational studies. Metagenomic shotgun sequencing has been shown to often provide high taxonomic and phylogenetic resolution [17, 18], but these approaches continue to be prohibitively expensive in many cases (particularly when in the presence of excess host DNA), and consensus remains in flux regarding the best pipelines for shotgun metagenomics-based community analysis [17].

An alternative is to use “3^rd^ generation” long-read sequencing technology to obtain full-length 16S rRNA gene sequences (V1-V9, hereafter FL16S). This increases taxonomic and phylogenetic resolution by increasing the number of informative sites sequenced, while continuing to use a well-studied pan-bacterial marker gene. Initial applications of Pacific Biosciences (PacBio) single-molecule real-time (SMRT) sequencing were hampered by the technology’s high intrinsic error rate [19-21], but improvements to the chemistry have since allowed for the generation of high-quality “circular consensus sequence” (CCS) reads, in which individual 16S rRNA genes are sequenced many times using circularized library templates combined with highly processive polymerases that provide for single-molecule, consensus-sequence error correction [22]. Recent studies evaluating FL16S sequencing by PacBio have found that, with appropriate processing and filtering, CCS reads of FL16S genes can be generated that are of sufficiently high quality to offer higher taxonomic resolution than partial 16S rRNA sequences [23-26].

The composition of the human sinonasal microbiome and how it changes in health and disease remains poorly understood, largely due to differences in methodology among studies resulting in large variations in reported bacterial profiles [27-32]. Culture-based approaches capture <15% of resident bacterial taxa when compared to nucleic acid-based techniques, since fast-growing bacteria like staphylococci tend to predominate in culture specimens, and recovery of anaerobes and slow-growing bacteria is limited [29, 33, 34]. Comparing across recent surveys of the sinonasal bacterial community reveals broadly similar results, but few specific assertions can be made; agreement between studies and results have been limited by an inability to distinguish bacteria at the species level [35-39] but as discussed above does not give a complete reflection of the microbial community. Thus, despite the vastly superior ability of molecular techniques to identify bacterial phylotypes, species-specific identification of bacteria remains superior in culture-based techniques [40]. For this reason, improved specificity of molecular detection techniques is necessary for not only a more complete understanding of the human sinonasal microbiome and other microbial communities, but also to be able to use this approach for decision making in the clinical context. Lastly, identifying the microbial taxa at play in different diseases with higher specificity will enable more directed experimental follow-up studies.

To take advantage of newer PacBio sequencing chemistry, improve upon data processing methods, and apply FL16S gene sequencing to a clinically relevant context, we describe a new pipeline (MCSMRT, “Microbiome Classification by Single Molecule Real-time Sequencing”). We show using two mock communities (one with 280 bacterial species) that FL16S CCS reads offer unprecedented accuracy and precision. We then explore bacterial diversity in the human nose and paranasal sinuses using results from MCSMRT, investigating not only bacterial diversity among subjects but also diversity within subjects at distinct sub-anatomical sites.

## Results

### Microbial community profiling by FL16S deep sequencing and CCS error correction

The taxonomic and phylogenetic resolution of microbial community profiling via 16S rRNA gene sequencing was increased by using Pacific Biosciences (PacBio) RSII to generate FL16S sequences from mock and human sinonasal microbial communities. We combined a circular sequencing template approach with the long DNA polymerase read-lengths provided by the PacBio sequencing technology. This provided for multiple sequencing passes of each molecule, enabling the generation of circular consensus sequence (CCS) reads of exceptionally high quality [20, 22]. To analyze these data, we developed a new bioinformatics pipeline, MCSMRT, building upon the uparse pipeline [41], which (a) processes and filters PacBio CCS reads generated from multiplexed samples, (b) *de novo* clusters high-quality FL16S sequences into “operational taxonomic units” (OTUs), (c) taxonomically classifies each read and assigns confidence values at each taxonomic level, and (d) quantifies the abundance of each OTU based on the full CCS read dataset (**Figure 1**). This processed data is suitable for downstream microbiome analyses using standard tools [42-45]. We further apply our classifier to all filtered reads and also allow for detection of amplicon sequence variants (ASVs) among groups of related sequences via minimum entropy decomposition (MED). Details are in the **Methods** and **Additional File 1**, and the MCSMRT software documentation which is freely available (https://github.com/jpearl01/mcsmrt).

**Figure 1.** Overview of the MCSMRT pipeline represented as a flowchart. The MCSMRT method for analysis of 16S rRNA reads from the PacBio is carried out in two steps: In the pre-clustering step, CCS reads are generated during demultiplexing, labeled by sample, pooled together, and then filtered based on several criteria (length distribution, terminal matches to the primer sequences, and not aligning to a host or background genome sequence). Before the clustering step, CCS reads are filtered based on cumulative expected error (EE<1). The clustering pipeline uses uclust to unique sequences based on their abundance, then cluster CCS reads into OTUs, filtering out chimeric reads during clustering, and then by using uchime after clustering. An OTU count table is created by mapping the filtered results from the end of the pre-clustering pipeline, and each OTU is taxonomically classified based on a representative “centroid” sequence. Taxonomic classification is also applied to all filtered reads, and ASV detection by MED can be applied on multiple alignments of sets of related sequencing, grouped by either OTU or binned by taxonomic level.

Below, we demonstrate the robustness and high taxonomic and phylogenetic resolution of our experimental and bioinformatics approach. We use results from two distinct mock microbial communities: one from the Biodefense and Emerging Infections Research Resource (BEI) and the other from the “Critical Assessment of Metagenome Interpretation” (CAMI) project[46]. We then applied FL16S gene sequencing to ask how the healthy human sinonasal microbiome varies among individuals and among sub-anatomical sites within individuals (**Table 1**; expected mock community compositions in **S1 Table** and **S2 Table, Additional File 2**).

**Table 1.** Community Characteristics*

|  | **BEI-EC** | **CAMI** | **HSNM-MS** |
| --- | --- | --- | --- |
| Source | Biodefense and Emerging Infections Research Resource Repository (BEI) | Joint Genome Institute (JGI) | Philadelphia Veterans Affairs Medical Center |

| Type | Mock Community | Mock Community | Wild Community |
| --- | --- | --- | --- |
| Details | BEI Even B (Catalog ID: HM-782D) | CAMI competition | Human Sinonasal Microbiome |
| Number of bacterial species | 20 | 282 (308 strains) | Unknown |
| Number of other species | 2 (archaeal and fungal) | 2 (archaeal) | Unknown |
| Species distribution | Reportedly even | Widely varying | Unknown |
| Other | Pooling DNA based on 16S qPCR | Pooling DNA based on genomic DNA mass | 12 subjects, 6 sites, swab and biopsy |

\*Source and composition characteristics of BEI-EC and CAMI (mock communities), and HSNM-MS (Human sinonasal community).

### CCS and filtering for FL16S reads

From each sample across the three types of communities, PCR was used to amplify FL16S genes (∼1.5 kilobases [kb]) from total DNA purifications using primers that targeted conserved regions at both ends of the gene that also contained terminal asymmetric barcodes to allow for pooling and subsequent demultiplexing of multiple samples into the same SMRTcell (**S3 Table, Additional File 2**). For the simpler BEI mock community, we tested PCR parameters, varying the polymerase (GoTaq vs. AccuPrime), cycle number (22 vs. 35), and the presence of excess off-target DNA (*i.e.* 10-fold excess of genomic DNA from U937 lymphoblast lung cell line).

We used the PacBio RSII (P6-C4 chemistry) to collect 3,446,849 polymerase reads in total across the 3 communities (**Table 2**). We typically obtained ∼50K-60K polymerase reads per RSII SMRTcell, in which 3-4 barcoded 16S amplimer libraries were pooled together and subsequently demultiplexed. More than half of polymerase reads were typically >20 kb, such that the average polymerase read included ∼12 complete sequencing passes around each molecule (average 1,422 base pair [bp] inserts). Those polymerase reads with >4 passes were used to generate error-corrected CCS reads, whose quality was dramatically improved (mean cumulative expected errors, EE, of 4.9 per kb) compared to polymerase reads to (EE / kb = 139.4). CCS reads with 1-4 passes had considerably lower quality (mean EE / kb = 193; 98.9% of these reads had EE > 10), and these were not considered further. The requirement for at least 5 passes resulted in a large reduction in overall yield compared to total polymerase reads but massively increased confidence in base calling (**Table 2**).

**Table 2.** Bulk sequencing and filtering stats for all three communities

| Source<sup>1</sup> | BEI-EC<sup>2</sup> | CAMI<sup>3</sup> | HSMC-MS<sup>4</sup> |
| --- | --- | --- | --- |
| Type | mock | mock | human |
| Total Samples | 8 | 1 | 122 |
| Total # SMRTcells | 4 | 1 | 85 |
| Total Pol Reads | 396,625 | 53,164 | 2,997,060 |
| N<sub>50</sub> Pol Read length in kb | 23523 | 21220 | 18837 |
| Avg Read Length in Pol Reads | 12997 | 11232 | 11940 |
| Avg Phred-quality in Pol Reads | 9.89 | 9.5 | 9.35 |
| Avg EE of Pol Reads<sup>5</sup> | 1868.82 | 1727.44 | 1653.9 |
| EE per kb of Pol Reads | 143.79 | 153.8 | 138.52 |
| Avg CCS passes | 14.04 | 13.06 | 11.48 |
| Avg Phred-quality in CCS Reads | 40.46 | 40.84 | 39.26 |
| Avg CCS length | 1454 | 1481 | 1417 |
| Avg EE for all CCS Reads<sup>5</sup> | 5.36 | 3.15 | 7.16 |
| EE per kb of CCS Reads | 3.69 | 2.13 | 5.05 |
| Total CCS Yield (>4 passes) | 163,689 | 19,576 | 787,302 |
| Size filtered (0.5-2kb) | 131,413 | 16,061 | 498,007 |
| Host filtered | 163,670 | 19,574 | 704,935 |
| Primer matched | 131,856 | 16,156 | 498,820 |
| Percent passed primary filters | 80.4 | 81.8 | 63.1 |

<sup>1</sup>In addition to mixed species communities, 7 independent negative reagent controls and 4 positive controls (2 x DNA from pure cultures of *Escherichia coli* and *Agrobacterium tumefaciens*) were run.
<sup>2</sup>Three conditions: Polymerase (GoTaq vs. AccuPrime) x PCR Cycles (35 vs. 22) x Excess DNA (no human DNA vs. 10-fold excess).
<sup>3</sup>Four independent libraries.
<sup>4</sup>Twelve subjects with healthy sinuses sampled at 6 sinonasal anatomic locations, both swab and biopsy. Twenty samples (mostly sites E and F) were not collected or not run.
<sup>5</sup>EE is cumulative expected error across the full read.

A series of additional filters were applied to these CCS reads to eliminate off-target sequences: (a) a size filter, (b) a filter against background (host) sequences, and (c) a primer matching filter. Collectively, these filters eliminated ∼20-40% of CCS reads (**Figure 1A**, **Table 2**, **S1 Figure, Additional File 3**, **S4 Table, Additional File 2**).

#### Size filter

CCS reads were removed if they were outside the thresholds of FL16S sequences (those between 0.5-2 kb were retained). This 2-3% of all CCS reads were mostly dimeric 16S sequences ∼3 kb long, most likely created during the ligation step of library preparation (**S2 Figure, Additional File 3**).

#### Host filter

CCS reads were removed if they aligned to a background genome (in this case, the human GRCh37, or hg19, reference). Notably, only 19 of ∼160K reads from the BEI mock community samples mapped to the human genome, despite half of these samples including a 10-fold excess human DNA (extracted from U937 human lymphoblast lung cell line to minimize contamination from the human microbiome). This indicates no appreciable off-target priming or contaminating fragments from U937 cell line DNA added at a 10:1 excess (by mass). However, samples with added human DNA had marginally lower sequencing yields (∼25%), possibly indicating a weak inhibitory effect by excess off-target DNA (**S3A Figure, Additional File 3**, effect on CCS yield, Tukey’s HSD p=0.048 for 10-excess human DNA, but p>0.2 for polymerase or cycle number).

By contrast, a much larger proportion of CCS reads from the human sinonasal communities mapped to the human genome (9.9%), suggesting off-target amplification of human DNA when in vast excess over bacterial DNA (alternatively, the U937 cell line DNA used for the BEI experiment may have lacked some or all off-target priming sites present in the human reference). Supporting this interpretation, biopsy samples had substantially higher total DNA yields after extraction than swabs (though no obvious differences in PCR yield, **S4 Figure, Additional File 3**), and reads derived from human were significantly more abundant in biopsy samples (**S3BC Figure, Additional File 3,** comparing biopsies and swabs, Tukey’s HSD p<<0.01 for total CCS yield or % human-contaminants, but p>0.8 when for patient or site). The 105,801 reads in the sinonasal dataset that mapped to the human genome aligned to 9,716 distinct genomic positions, but they were highly enriched at only a few (67.9% of mapped positions had only a single mapped read, but 58.2% of reads started at only 16 positions and had >1,000-fold coverage, **S5 Figure, Additional File 3**). These data suggest an off-target priming effect at high excess concentrations of human DNA with “hotspots” for off-target priming, along with a proportion of library molecules carrying apparently random human genomic DNA fragments. To confirm that reads mapped to the human genome were not improperly aligning true bacterial 16S genes, utax classification of all human-mapping reads showed that all had extremely low confidence assignments to the bacterial domain (<0.1), indicating a probable host origin.

#### Primer matching

CCS reads were required to have the forward and reverse primer sequences each found once and oriented correctly at the ends of the sequence, and this removed 12-18% of reads (**Table 2, S4 Table** and **S13 Table**, **Additional File 2, S6 Figure, Additional File 3, Additional File 1**). Primer matching also served to determine the orientation of the 16S gene in each CCS read, so reads were reverse complemented when the reverse primer came first. Finally, primers were trimmed from reads. In principle, this loses several taxonomically informative sites, since the primers contained four degenerate bases; however, in practice, the primer sequence seen in a given read was random with respect to the taxonomic source of that 16S gene. This is most easily illustrated from control sequencing of 16S rRNA genes amplified from clonal cultures of *Escherichia coli* K12 MG1655 and *Agrobacterium tumefaciens* NTL1 (**S7 Figure and S8 Figure, Additional File 3**).

### Clustering CCS reads into OTU

Profiling the bacterial composition of a microbiome often begins by clustering sequences with high sequence identity into OTU, with a standard cutoff of 97% [47, 48], though sometimes other cut-offs are used [49-51]. Newer approaches to grouping related sequences together avoid using similarity thresholds but instead define amplicon sequence variants (ASVs) based on controlling for variant sites arising due to sequencing error; these methods include oligotyping, minimum entropy decomposition (MED), and DADA2 [44, 45, 52]. Here, we initially show results with OTU clustering and then further show how MED can further discriminate species whose 16S rRNA genes diverge by less than the threshold used for OTU picking.

To first cluster reads and identify representative “centroid” OTU sequences, we used the UCLUST algorithm [53](**Additional File 1**), which filters chimeric 16S sequences by identifying apparent hybrids between distinct OTU as they accumulate in the dataset (here called CHIM1). A second chimera filter (CHIM2, using uchime) [54] then removes centroid OTU that appear to be hybrids of distinct 16S sequences in the curated Ribosomal Database Project (RDP) Gold database [55] (**Additional File 1**). The abundance of each OTU in each sample was then determined by counting the number of filtered CCS reads that aligned to each centroid (**Figure 1B**). Though OTU clustering can collapse or separate distinctly named species into the same or different OTU, it systematically defines taxa in a uniform way that does not depend on taxonomic nomenclature [56, 57].

#### Sequencing error increases observed OTU counts

Erroneous base calls in CCS reads risk artificially inflating the number of OTUs, since reads with sequencing errors in similar 16S rRNA genes can be spuriously separated into distinct OTUs, especially if their actual divergence is near the 3% divergence cut-off and/or they are short. Thus, a final pre-clustering filter was applied based on cumulative expected error (EE, or the sum of error probabilities across all positions in a read as determined from Phred-scaled base quality scores). This measure has previously been shown to discriminate against error prone sequences better than the average quality score [58].

#### Analysis of the BEI mock community

Using the BEI mock community to examine the relationship between CCS passes and EE, we found, as expected, that reads with more CCS passes had a lower median EE (**Figure 2**, linear model of log(EE) vs. CCS passes gives R^2^=0.22). 98.9% of reads with less than five CCS passes had EE>10 (across ∼1.4 kb total length) and were not considered further (**Figure 2**).

**Figure 2.** Distribution of reads at different CCS passes and cumulative expected error values (EE) in the BEI mock community. Violin plot showing the distribution of cumulative EE (after primer matching and trimming) at different CCS passes. Reads with less than two CCS passes were not reported by PacBio CCS software. Histograms at the top and right show read count by CCS and EE, respectively. The 35 reads with 26 to 46 CCS passes are not show (median EE = 0.22). Subsequent analyses used only CCS reads with >4 passes.

To empirically determine an appropriate EE cut-off for clustering CCS reads into OTUs, we compared the expected number of OTUs in the BEI mock community to that obtained by OTU clustering at different EE cut-offs. The expected number of OTUs was 19, since 2 of the 20 species’ 16S rRNA genes differed by at most 23 nucleotides (*Staphylococcus aureus* and *Staphylococcus epidermidis* have only 1.4% divergence, less than the OTU clustering cut-off of 3% divergence, **S1 Table, Additional File 2**). As expected, increasing the stringency of the EE filter reduced the total number of CCS reads available for OTU clustering, as well as the total number of OTUs detected (**Figure 3A**). Using EE≤1 (one or fewer expected errors per read) retained less than half (40.1%) of filtered CCS reads, but these clustered into the 19 OTUs expected. Decreased stringency (higher EE cut-offs) increased the total OTUs detected, dramatically for cut-offs of EE ≤8 and above; using no expected error threshold (EE ≤128), 3,453 OTUs were detected, more than 100-fold greater than the true number. In summary, using a high stringency expected error cut-off for OTU clustering reduced the number of reads available for clustering but provided exact total OTU counts for the BEI mock community.

**Figure 3.** Clustering of post-filtered CCS reads into OTUs. **A**. Count of total, unique, CHIM1 and centroid OTU reads at different maximum EE thresholds. **B**. Count of total OTU detected using full-length or truncated reads at different maximum EE thresholds.

#### Sequence length and OTU clustering

To examine how OTU clustering would be affected by using partial instead of FL16S gene sequences for the BEI mock community, we performed *in silico* primer matching and trimming on the full-length CCS reads for three short-read primer pairs commonly used for microbial community profiling, namely primers targeting the V1-V3, V3-V5, or V4 hypervariable regions of the 16S rRNA gene, which are amenable to analysis using Illumina short-read sequencing (**S3 Table, Additional File 2**). Using these *in silico* short-read data produced dramatically higher OTU counts than predicted, even when using substantially more stringent EE cut-offs (**Figure 3B**). Thus, for example, whereas full-length V1-V9 reads clustered into the expected 19 OTUs at EE ≤1, reads truncated to include only the V3-V5 hypervariable regions (average = 536 nt) clustered into 116 OTUs. The other truncated sequences (V1-V3 and V4) had even higher elevated total OTU counts. Even when the shorter length of partial sequences was compensated for by using an 8-fold higher stringency cut-off (EE ≤ 0.125), spurious OTUs were still detected, e.g. 58 OTUs were detected with the V3-V5 truncated reads, substantially higher than expected. The comparisons above relied on *in silico* truncation of full-length reads from the same PacBio dataset to maintain consistent error profiles, but inflated OTU counts have also been reported in published results for truncated 16S from the BEI mock community collected using both 454 pyrosequencing and Illumina MiSeq short-read technology [59-63]. For comparison, we also applied closed-reference OTU clustering at a 97% cutoff via QIIME2, and found inflated OTU counts using either FL16S (115 OTUs) or V3-V5 truncated reads (202 OTUs) (if counting only OTUs with >4 reads mapping, then FL16S detected 60 OTUs and V3-V5 detected 115 OTUs), though FL16S still detected fewer OTU counts than truncated reads. Finally, we similarly found elevated OTU counts in a re-analysis of Illumina MiSeq data for the V1-V3 region of the same BEI mock community [62] through our pipeline, finding 171 OTUs at EE<1 (40 OTUs at EE<0.125). This suggests that inflated OTU counts when using partial 16S sequences is independent of the specific PCR conditions or the particular error profile of PacBio CCS reads.

The above results underline the value of using FL16S to minimize the effect of sequencing errors on *de novo* OTU cluster counts. They also indicate that methods that profile taxonomic composition using partial 16S rRNA genes may be prone to overestimating bacterial diversity. For all subsequent analyses, we used only CCS reads with EE ≤1 for OTU clustering, and then mapped all reads passing all pre-clustering filters onto these centroid OTU’s to obtain abundance data (**Pre-clustering Pipeline, Clustering Pipeline, Additional File 1)**.

### Taxonomic classification of FL16S reads

Bacterial taxonomic nomenclature has traditionally been based on physiological and other microbiological traits (*e.g.* virulence) rather than 16S rRNA gene sequences, so the accuracy and precision with which a read can be taxonomically defined is dictated by: a combination of organism-specific criteria used for naming species; the quality and completeness of the database used; and the distribution of informative variable sites within the 16S rRNA among named taxa [10]. Unfortunately, commonly used databases for classifying 16S rRNA gene sequences, namely RDP and Silva [55, 64, 65] (although see [66] for a new way of extracting FL16S reads with species level classifications from RDP), do not provide species-level taxonomic identifiers [67]. Another problem with these databases is the absence of representative sequences from genera present in our mock communities (for example the genus *Clostridium* was not found in RDP). Though 285,289 sequences in the popular Greengenes database do have species labels, only 631 of these are unique species. Although Greengenes (N=1,262,986) and Silva (N=1,922,213) have vastly more taxonomically classified sequences than RDP (N=8,978), in part because they computationally assign taxonomies to sequences from environmental microbiome surveys [64, 68], most of the sequences in these databases are of partial length 16S rRNA genes. While these databases are appropriate in many cases, we needed to make a database of FL16S sequences with species-level taxonomic information.

To classify CCS reads (including centroid OTUs and MED node representatives) based on bacterial taxonomy to the species level and also provide confidence values at each taxonomic level, we trained a utax classifier on a custom-built database of FL16S gene sequences downloaded from NCBI (**16S rRNA Microbial Database, Additional File 1**). Most FL16S sequences available at NCBI (N=17,764) could be associated with a taxonomic ID (txid) using the gid accession number, allowing us to extract, parse, and configure sequences in the database to create a utax-compatible Linnaean hierarchy that included 11,055 distinctly named species spanning 367 bacterial families (**Additional File 1**). The number of distinct families present in this NCBI database was 367, whereas RDP had 366, Silva had 302, and Greengenes had 514. We recognize that other researchers may not prioritize species-level taxonomic assignments and instead favor high breadth. To that end—because MCSMRT is based on the UPARSE pipeline—any correctly formatted database may be used in place of our custom one. UPARSE-formatted databases for Greengenes, Silva, and RDP are available and may be found at [69].

We next generated and compared the accuracy of utax classifiers built from full-length or partial V3-V5 16S rRNA genes by classifying the database sequences themselves. In this context, incorrect classification could arise in particular due to distinct named species with highly similar sequences. The full-length classifier gave an incorrect label only 1.0% of the time (N=173 mistakes), compared to 13.2% of the time using the truncated classifier (N=2295 mistakes). Indeed, when the two classifiers disagreed, the full-length call was much more frequently correct (2x2 contingency table: 15,081 both correct, 2,137 only full-length correct, 15 only truncated correct, and 158 neither correct). Furthermore, species-level confidence values were higher 81.3% of the time using the full-length classifier (mean 81.7%, median 92.7%) compared to the truncated classifier (mean 71.1%, median 82.7%). These results show the value of using full-length compared to partial 16S gene sequences for accurate taxonomic assignment.

The assignment made for each CCS read is associated with confidence values at each taxonomic level, and low values could arise for several reasons aside from the quality of the sequence data. In particular, sequences labeled as a distinctly named species could have other equally good matches, or nearly so. In order to determine what species might end up assigned to a particular centroid OTU read, we clustered the NCBI database sequences (17,776 in total, 99.1% unique) at the same threshold level (97% identity), thereby grouping species belonging to the same “database OTU” (dbOTU). Since uclust relies on abundant unique sequences to initiate centroids and also drops putative chimeric sequences during clustering, we instead applied hierarchical clustering (average linkage, using pairwise percent identity values from all-by-all blast, and separating dbOTU clusters at a 3% difference level). This method is unaffected by the order of the sequences and included all database entries.

Hierarchical clustering of NCBI sequences resulted in 6,065 dbOTU, of which 66.9% of clusters had a single species (93.2% had a single genus), whereas 14.6% of clusters had the same species split over more than one dbOTU (**Additional File 4**). Some dbOTUs consisted of many species. For example, the top three most species-rich dbOTUs collectively contained 453 distinctly named *Streptomyces* species indicating that 16S rRNA clustering at 3% divergence poorly discriminates among named species in this genus [70]. These results reflect the variability with which different bacterial taxa are named compared to how they group based on divergence in their 16S rRNA [50, 51] (**S9 Figure, Additional File 3**). Collectively, clustering the FL16S gene sequences from NCBI indicated that assignments of individual CCS reads, OTU centroids, or MED node representatives will result in high-confidence species-level classification for high quality FL16S gene sequences; imprecision due to distinct species belonging to the same dbOTU will affect about a third of 16S rRNA sequences in the database, but these can be flagged by low confidence values from the utax classifier and by cross-referencing to dbOTU clusters to identify other possible “nearly best hits”.

**Nearly all reads collected from reagent controls failed filtering and classification steps.** In advance of the studies described above, we pre-screened multiple PCR reagents and DNA polymerases to identify those that produced no observable amplification when using reagent controls. In addition, seven negative control reagent samples were sequenced in parallel with mock community and sinonasal samples. After demultiplexing, these controls produced a total of only 54 reads. Of these, only 3 reads passed the filtering criteria described above. Taxonomic classification on all 54 reads returned only 5 reads with >10% bacterial domain-level confidence values (all 5 gave 100% confidence). These 5 reads had genus-level classification as *Finegoldia* (95.3% confidence), *Propionibacterium* (16.6%), *Streptococcus* (22.7%) and two as *Staphylococcus* (0.8%). Four of these reads had 0% species-level confidence, while *Finegoldia magna* had 94% species-level confidence. These results demonstrate that our laboratory and bioinformatics methods produce extremely low levels of contamination from off-target bacterial nucleic acids generated from our reagents. Further consideration of controlling for contaminants would require direct empirical measurements of 16S copy number in each sample [71].

### BEI mock community composition

#### OTU classification

The 19 distinguishable OTUs in the BEI mock community were readily identified and accurately classified (**Table 3**). All but 3 of the centroid OTU— including 3 distinct *Streptococcus* species—were correctly classified to the species level. The 3 discrepancies were, however, reflected by low confidence values assigned by utax, as well as by clustering of discrepant taxa into the same dbOTU.

**Table 3.** Taxonomic classification of BEI*

| Centroids Full-Length |  |  | Centroids v3-v5 |  |  | Clustered from v3-v5 truncated reads |  |  |
| --- | --- | --- | --- | --- | --- | --- | --- | --- |
| Genus_Species | Genus Conf | Species Conf | Genus_Species | Genus Conf | Species Conf | Genus_Species | Genus Conf | Species Conf |
| *A. baumannii* | 0.9796 | 0.9392 | *A. baumannii* | 0.8784 | 0.4225 | *A. baumannii* | 0.8784 | 0.4225 |
| *A. odontolyticus* | 0.9897 | 0.9392 | *A. odontolyticus* | 0.9728 | 0.6189 | *A. odontolyticus* | 0.9728 | 0.6189 |
| *B. anthracis* | 0.9876 | 0.494 | *B. cereus* | 0.963 | 0.4225 | *B. cereus* | 0.963 | 0.3114 |
| *B. vulgatus* | 0.993 | 0.9438 | *B. vulgatus* | 0.9847 | 0.897 | *B. vulgatus* | 0.9847 | 0.897 |
| *C. beijerinckii* | 0.989 | 0.7194 | *C. roseum* | 0.9669 | 0.4225 | *C. roseum* | 0.965 | 0.3114 |
| *D. radiodurans* | 0.9935 | 0.9438 | *D. radiodurans* | 1 | 0.897 | *D. radiodurans* | 0.9994 | 0.897 |
| *E. faecalis* | 0.9862 | 0.9392 | *E. faecalis* | 0.8784 | 0.8265 | *E. faecalis* | 0.8784 | 0.8265 |
| *S. flexneri* | 0.6878 | 0.494 | *S. flexneri* | 0.6661 | 0.4225 | *S. flexneri* | 0.6072 | 0.3114 |
| *H. pylori* | 0.9918 | 0.9346 | *H. pylori* | 0.9787 | 0.9022 | *H. pylori* | 0.9787 | 0.9022 |
| *L. gasseri* | 0.9928 | 0.7194 | *L. gasseri* | 0.9728 | 0.4225 | *L. gasseri* | 0.9728 | 0.4225 |
| *L. monocytogenes* | 0.989 | 0.7194 | *L. ivanovii* | 0.961 | 0.4225 | *L. ivanovii* | 0.961 | 0.3114 |
| *N. meningitidis* | 0.973 | 0.9346 | *N. meningitidis* | 0.9533 | 0.6189 | *N. meningitidis* | 0.9533 | 0.6189 |
| *P. acnes* | 0.973 | 0.9575 | *P. acnes* | 0.9669 | 0.9099 | *P. acnes* | 0.965 | 0.9099 |
| *P. aeruginosa* | 0.973 | 0.9346 | *P. aeruginosa* | 0.9329 | 0.6189 | *P. aeruginosa* | 0.9329 | 0.6189 |
| *R. sphaeroides* | 0.973 | 0.5993 | *R. sphaeroides* | 0.9228 | 0.4225 | *R. sphaeroides* | 0.9228 | 0.3114 |
| NA | NA | NA | NA | NA | NA | *S. aureus* | 0.1509 | 0.0089 |
| *S. epidermidis* | 0.9869 | 0.7194 | *S. epidermidis* | 0.6661 | 0.4225 | *S. epidermidis* | 0.6661 | 0.3114 |
| *S. agalactiae* | 0.9897 | 0.9483 | *S. agalactiae* | 0.9728 | 0.8996 | *S. agalactiae* | 0.9728 | 0.8996 |
| *S. mutans* | 0.9918 | 0.9529 | *S. mutans* | 0.9768 | 0.8996 | *S. mutans* | 0.9768 | 0.8996 |
| *S. pneumoniae* | 0.9904 | 0.494 | *S. pneumoniae* | 0.9768 | 0.4225 | *S. pneumoniae* | 0.9768 | 0.4225 |

\*Classification results from the BEI mock community. The performance of FL16S centroid OTU assignments compared to those truncated to V3-V5. The last two columns indicate the top 20 most abundant OTU from truncating to V3-V5 prior to clustering.

All 19 centroid OTUs were correctly classified to the family level, with one genus-level discrepancy: classification of *Escherichia coli* as *Shigella flexneri*. This incorrect assignment is not a surprise; indeed, the matching dbOTU contained 54 sequences that were assigned to *Escherichia*, *Shigella*, *Citrobacter*, and *Salmonella*, all genera known to have low levels of divergence among their 16S rRNA genes[72, 73]. Two species-level assignments were incorrect: (a) *Bacillus cereus* was classified as *Bacillus anthracis*; these two differ by only 2 nucleotides in their 16S genes and share the same dbOTU with 8 other *Bacillus* species; and (b) *S. aureus* and *S. epidermidis*—whose 16S genes differ by 23 nucleotides—were collapsed into a single OTU, with the centroid called *S. epidermidis* (the matched dbOTU consisted of 39 additional staphylococci, though see below). Species-level confidence values were 0.49 and 0.71 respectively. By contrast, only one correctly classified OTU, *Streptococcus pneumoniae*, had a species-level confidence value <0.50 and this species shared the same dbOTU with 13 additional *Streptococcus* species.

Truncation of CCS reads prior to clustering considerably worsened classification; in addition to increasing the number of total OTU, CCS reads truncated to their V3-V5 region prior to clustering resulted in seven misclassified OTU among the top 20 most abundant OTU (**Table 3**). To isolate the effects of truncation on classification alone, rather than both clustering and classification, we also truncated the 19 centroid OTU to their V3-V5 region and classified these using a utax classifier built from a database of sequences also truncated to V3-V5. This showed reduced species-level confidence values but also more miscalled taxa (**Table 3**). These results show that use of FL16S gene sequences provides substantially improved taxonomic identification of centroid OTU compared to truncated 16S rRNA sequences, to the extent that bacterial nomenclature allows.

For comparison, we used the Greengenes v13_8 database to classify closed-reference OTU identified by QIIME2. The resulting classifications were typically at higher taxonomic levels and more often incorrect. For example, using V3-V5 sequence, 5 of the top 19 most abundant closed-reference OTUs had species level annotation (*S. agalactiae*, *L. seeligeri*, *C. paraputrificum*, *S. saprophyticus*), but only one of these matched a species in the BEI community (*S. agalactiae*). Greengenes classification with FL16S made only two classifications to the species level, both incorrect (*S. saprophyticus* and *Alkanindiges illinoisensis*). Often classifications were to much higher levels (*e.g.* family Planococcaceae or even the domain bacteria). Taxonomy results using QIIME2 are reported in **S5 Table, S6 Table, S7 Table,** and **S8 Table** in **Additional file 2**.

#### Relative abundance and sequencing error

The abundance of each OTU was estimated by assigning all filtered CCS reads (with no EE threshold) to a centroid OTU with a maximum of 3% divergence for a hit to be counted. The twenty bacterial species in the BEI mock community were expected to have equimolar abundances of their 16S rRNA genes, and for most species, we detected a roughly even mock community composition for most species (**S10 Figure, Additional File 3**). Several were outliers: (a) the two *Staphylococcus* species were binned together as *S. epidermidis* (as described above); (b) *Bacteroides vulgatus* and *Helicobacter pylori* were overrepresented, especially at high PCR cycle number; and (c) five species were found at lower than expected abundances across PCR conditions.

We next evaluated the impact of (a) chimeric sequences on relative abundance measurements and (b) “true” substitution errors. First, all CCS reads were run through uclust with no filters other than requiring >4 CCS passes to identify likely CHIM1 chimeras, and then all CCS reads were aligned to the 16S reference sequences from the BEI community to determine their likely source. This again found relatively even abundances for each taxon with the exception of those mentioned above. Increased cycle number also increased the variance among taxa in their relative abundances, but the inclusion of chimeric reads had little effect (**S11 Figure, Additional File 3**).

Second, the number of base mismatches (excluding gap characters) was calculated between each non-chimeric read and its most similar reference sequence, estimating the number of “true” substitution errors made during sequencing (though intragenomic variation in 16S rRNA gene sequences [74] also contributes to putative substitution errors). This analysis indicates that the AccuPrime polymerase made fewer errors than GoTaq polymerase but that errors made by either polymerase were insufficient to inflate OTU numbers when using full-length sequence (**S12 Figure, Additional File 3**). Initial analysis of the number of errors from the *E. coli* reads suggested that the single reference FL16S copy may not have been correct (almost no reads were an exact match) (**S12A Figure**).

To further examine sequencing error, we investigated sequence variation in FL16S CCS reads collected from our *E. coli* K12 MG1655 monoculture positive control samples. We first obtained a finished circular assembly of our lab’s strain by shotgun sequencing on the PacBio RSII, and we identified and extracted seven FL16S genes (two were identical, but the other all differed slightly). This allowed us to obtain more confident estimates of the true error rate in individual CCS reads by globally aligning all 8028 reads to their closest matching FL16S copy from the genome (**S13 Figure, Additional File 3)**. Reads with EE≤1 had considerably lower “true” error rates (mean mismatches = 3.0, mean gaps =0.8), compared to those with EE>1 (mean mismatches = 5.5, mean gaps = 6.0), illustrating an especially dramatic loss of error due to gaps after EE filtering. Additionally, we used MED to identify ASVs from multiple alignments of *E. coli* positive controls. The resulting MED node representatives were aligned with the 16S genes identified from our whole genome assembly, and an approximate ML tree shows that ASVs correctly segregated with individual genomic 16S copies (**S13B Figure, Additional File 3**).

Independent analyses of the same BEI mock community by Illumina MiSeq for V3-V5 have previously found the same taxa elevated or depleted, suggesting that these taxa actually are at unequal concentrations in this mock community [75, 76]. The primers we used have perfect identity with all BEI bacterial strains’ reference 16S rRNA gene sequences and are distinct from the Illumina-based analyses, so the compositional biases seen are not likely to be due to primer choice or PCR conditions [62].

#### Discriminating among closely related sequences

Because using FL16S should increase the number of taxonomically and phylogenetically informative sites, we reasoned that species whose 16S genes differ by less than the OTU clustering threshold would be more easily separated with full-length versus truncated 16S gene sequences. Although the two clinically important *Staphylococcus* species in the BEI mock community belonged to the same dbOTU (along with 40 other staphylococcal species, and 2 additional genera) and were not separated during *de novo* OTU clustering, they were readily distinguishable in several ways:

First, direct classification of primer-matched CCS reads from the BEI mock community identified only *S. aureus* and *S epidermidis* among those classified to the staphylococci (1649 from *S. aureus* and 2501 from *S. epidermidis*; using unfiltered CCS reads yielded 0.71% classified to 5 additional staphylococcal species in 32 reads). Thus, direct taxonomic classification correctly identified both species in roughly equal proportions.

Second, we applied MED to identify ASVs for all primer-match CCS reads with EE≤1 that had been assigned to the *Staphylococcus* OTU, and the MED node representatives were taxonomically classified. MED reduced 3171 CCS reads to 48 nodes (ASVs), and representative sequences from each ASV were used to build phylogenetic trees, also including all NCBI entries for the staphylococci (**Figure 4**). The results with FL16S sequences show that *S. epidermidis* and *S. aureus* and clearly separated from the other staphylococci, as expected, and node representatives classified as each species formed clear monophyletic groups (**Figure 4A**). By contrast, building trees from ASVs identified from the V3-V5 truncated reads did not clearly distinguish among staphylococcal species, and although in this case the node representatives were correctly classified, NCBI entries of other species were intermixed with those of the two expected species (**Figure 4B**).

**Figure 4.** Approximate maximum likelihood phylogenetic tree reconstruction of staphylococcal 16S sequences representing the ASV nodes identified by MED, along with staphylococcal NCBI database entries (midpoint rooting). Each filled tip symbol represents a single MED node, and its size represents the number of reads belonging to that node. Unfilled symbols indicate NCBI database entries. Color indicates the taxonomic assignment for the two expected species with others indicated with grey. **A.** Using FL16S (48 MED node representatives) **B.** Using truncated V3-V5 16S (33 MED node representatives).

These results show that distinguishing among closely related organisms—even when sequence differences are insufficient to separate these into distinct OTUs—is strongly facilitated by use of FL16S gene sequences, and especially powerful when combined with an ASV detection method.

### CAMI mock community composition

#### OTU classification

Because a curated set of FL16S gene sequences was not available for the 280 unique bacterial species present in the CAMI mock community, we first cross-referenced the the expected bacterial composition (**S2 Table, Additional File 2**) with the FL16S gene sequences in the NCBI database, finding one or more full-length sequences for all but 3 species, for which a taxonomy was available but not a corresponding 16S rRNA gene sequence (*Mameliella alba*, *Fusobacterium naviforme* and *Promicromonospora flava*). In addition, 11 species names used by CAMI and NCBI were synonyms, due to revisions in species names (particularly members of the *Clostridiales* family, **S9 Table, Additional File 2**). By cross-referencing the expected species with the NCBI dbOTUs, we found the 280 CAMI species would cluster into 253 OTUs at the 3% divergence level, each associated with a distinct dbOTU (except the three missing species). For example, 5 species from the genus *Prauserella* (*aidingensis, alba, flava, halophila, salsuginis*) existed in the CAMI community, and although there was an instance of each of those species in the NCBI database, none of these NCBI 16S sequences differed from each other by greater than 3%. Therefore, the expectation was a single OTU associated with the genus *Prauserella*, which was indeed the result. Most of these expected clusters had three or fewer named species in their corresponding dbOTU (84.4% of CAMI clusters), but they collectively comprised 586 distinct species calls in the NCBI database. This clustering allowed us to cross-reference the centroid OTUs to members of the CAMI community and identify ambiguities in the extant taxonomic classification (**S9 Table, Additional File 2**).

FL16S gene sequencing by PacBio had exceptionally high specificity and sensitivity for identifying the bacterial constituents within the complex CAMI mock community (**Figure 5**). The ∼16K filtered CCS reads (final yield from one PacBio SMRTcell) clustered into 227 OTUs (using 6,878 reads at EE ≤1) with 216 unique species names. Of these, 192 centroid assignments perfectly matched up with an expected cluster in the CAMI community, thus giving 89% exact species-level matches with the centroid OTU. Nineteen more centroids could be connected to CAMI-defined clusters, either via a dbOTU (13 centroids) or a correct genus-level assignment (6 centroids).

**Figure 5.** CAMI mock community composition. Observed count versus expected relative abundance, based on matching centroid OTU assignments with expected species composition.

Of the 5 remaining OTUs detected by PacBio that did not belong to a CAMI cluster, 2 could be accounted for with family-level matches to CAMI clusters (the Rhodobacteraceae *Mameliella alba*, absent from the NCBI database, was classified as *Paracoccus versutus*, and *Promicromonospora flava* was identified as *Isoptericola variabilis*). This left only 3 “false positive” OTUs, which accounted for a total of 7 CCS reads. Two of these misidentifications were species belonging to families represented in the CAMI community—the Rhizobiaceaen *Agrobacterium larrymoorei* (1 CCS read) and the Lachnospiraceaen *Moryella indoligenes* (5 CCS reads)—and one was not represented (the Moraxellaeceaen *Acinetobacter septicus* had a single CCS read).

Forty expected CAMI clusters were not identified among the centroid OTU (*i.e.* “false negatives”). This was, at least in part, due to under-sampling: The relative abundance of 16S rRNA genes for most CAMI species was expected to be very low (some well below our limit of detection), and all missing CAMI members had expected abundances of <1% (**Figure 5**). Another potential reason for failure to detect specific taxa would be failed amplification due to variation from our universal primers from CAMI community members. Since we derived four independent sequencing libraries from the CAMI community, each library effectively represented an experimental rarefaction, allowing us to ask how sensitivity was affected in single libraries compared to the total. On average, 20 OTUs were missing from individual libraries (with an average of 3,822 reads each) that had been captured when the four datasets were combined. Thus, with a single SMRTcell, we detected 84.2% of taxa present with 95% species-perfect identification, and nearly every single OTU identified by FL16S gene sequencing could be attributed to a member of the CAMI community. By contrast, truncating CCS reads (or their centroid OTUs) to the V3-V5 region was less accurate and showed lower species-level confidence values, as seen above for the database sequences themselves (see above and **S10 Table, Additional File 2**).

#### Relative abundances in the CAMI mock community

The expected relative abundance of each species in the CAMI mock community was accurately reflected by the number of reads assigned to each CAMI centroid OTU by usearch (**Figure 5**, **S9 Table, Additional File 2**). The expected relative concentration of each species’ 16S rRNA genes in the CAMI community was calculated using: (a) the genome size estimated by from *de novo* assembly of shotgun sequence collected from each CAMI accession [77] (**S9 Table, Additional File 2**), and (b) an estimate of 16S rRNA gene copy number using rrnDB [78]. For CAMI species missing from the rrnDB, the lowest Linnaean rank with members of the database was determined, and the average 16S rRNA copy number of all species under that rank was used (**S9 Table, Additional File 2**). Remarkably, we observed a strong linear fit between observed and expected abundances (**Figure 5**, R^2^ = 0.63), showing that we not only accurately identified the species present by centroid OTU, but also accurately quantified their relative abundances, despite the low expected relative abundance of most species’ 16S rRNA genes.

#### Phylogenetic discrimination of species in the same genus within the CAMI community

Because the CAMI mock community included 45 multi-species genera (9 with >3 species), we next asked whether FL16S reads discriminated among species in the same genus better than truncated V3-V5 reads. We collected all filtered CCS reads that had been classified to a given multi-species genus and produced phylogenetic trees from multiple sequence alignments of each genus-specific read set (39 multi-species genera with at least 5 filtered CCS reads).

Using these genus-level trees, we next assessed whether the utax-assigned species labels for each read formed monophyletic clades using MonoPhy [79] (**S11 Table, Additional File 2**). For most genera—where the species were sufficiently diverged— trees built from either FL16S or V3-V5 truncated reads performed comparably: For 28 of 39 genera, all assigned species labels were monophyletic using either FL16S or V3-V5 reads. Examples of well-resolved genera with either marker gene length included *Clostridium* and *Desulfovibrio* (**S14 Figure, Additional File 3**). Five more genera were non-monophyletic for an equal number of species using either marker gene length; some of this is likely due to poorly resolved species nomenclature. Examples include the genera *Azotobacter* and *Nonlabens* (**S15 Figure, Additional File 3**). For the remaining six multi-species genera, phylogenies built from FL16S reads showed higher monophyletic grouping of species-level classifications than trees built from V3-V5 truncated reads. Two prominent examples were closely related species within the *Algoriphagus* and *Salegentibacter* (**S16 Figure, Additional File 3**). In *Algorophagus*, *A. yeojeoni* appears polyphyletic for V3-V5 only, and in *Salegentibacter*, *S. salegens* fails to resolve from *S. salinarum*. These results further demonstrate the utility of increasing the length of marker gene sequencing to capture more informative sites, thus improving phylogenetic resolution of distinct but closely-related members of microbial communities.

### The composition of the human sinonasal bacterial microbiome

Rhinosinusitis effects 16% of the US population [80] and accounts for 1 in 5 antibiotic prescriptions to adults in the US in the outpatient setting, making it the most common diagnosis for outpatient antibiotic use in the U.S. [81]. Thus, a more complete understanding of the resident microbial community of the upper respiratory track is paramount to improved therapeutic interventions and reduction of inappropriate antibiotic prescriptions. Thus, we applied our bacterial microbiome profiling method to the human sinonasal cavity. We obtained samples from 12 subjects undergoing pituitary gland adenoma removal, utilizing the sinonasal cavity as a surgical corridor for access to the gland. None of the total 12 patients examined had objective or subjective findings of infectious or inflammatory disorders of their sinonasal complex. In creating a surgical corridor for access to the skull base, 6 distinct anatomical locations within the sinonasal cavity were sampled by both swab and biopsy (**Figure 6, Table 4, S12 Table, Additional File 2)**.

**Table 4.** Codes used for Swab/Biopsy sites

| Swabs |  | Biopsies |  |
| --- | --- | --- | --- |
| Code | Site | Code | Site |
| A | Nasal Vestibule | B | Head of Inferior Turbinate Tissue |
| C | Middle Meatus | D | Uncinate Process Tissue |
| E | Maxillary Sinus | F | Maxillary Sinus Tissue |
| G | Ethmoid Culture (Deep to Ethmoid Bulla) | H | Ethmoid Tissue (Deep to Ethmoid Bulla) |
| I | Superior Meatus | J | Sphenoethmoidal Recess Tissue |
| K | Sphenoid | L | Sphenoid Tissue |

**Figure 6.** Schematic diagrams in the sagittal and coronal planes of the human sinonasal cavity. Surgical access to the sella turcica (pituitary) is shown by the shaded arrow. Sites of sampling for microbiome analysis: deep nasal vestibule swab, deep to the vibrissae past the squamous mucosal epithelial junction (A), head of inferior turbinate swab (B), middle meatus swab (C), uncinate process biopsy (D), maxillary sinus swab (E) and biopsy (F), ethmoid sinus swab (G) and biopsy (H), superior meatus swab (I) and biopsy (J), and sphenoid sinus swab (K) and biopsy (L). Figure adapted from “Atlas of Endoscopic Sinus and Skull Base Surgery,” ed. Palmer, J.N., Chiu, A.G., Adappa N.D. Elsevier, Philadelphia (2013).

To determine the bacterial constituents of the human sinonasal microbiome and the extent to which it varies among healthy individuals and among distinct sinonasal sites, we sequenced FL16S amplicons by PacBio from paired swabs and biopsies at the 6 anatomical sites from the 12 subjects (**S12 Table, Additional File 2**, 122 specimens total across 12 individuals). No filtered CCS reads were generated from 23 samples, primarily from the maxillary sinus (both swabs and biopsies), suggesting little colonization of this site by bacteria (**Figure 6**), and significantly fewer reads were collected from biopsy samples than from swab samples, potentially indicating lower overall bacterial load compared with the mucosal surface (**S3 Figure, Additional File 3**). Filtering, clustering at an EE ≤1, taxonomic assignment, and counts per OTU per sample were conducted as above (counting all filtered CCS reads against all non-chimera centroid OTU sequences via usearch). Complete information about counts per OTU per sample, as well as the taxonomic assignments of each centroid OTU are in **S13 Table, S14 Table** and **S15 Table, Additional File 2,** and the results for all three communities have been incorporated into individual phyloseq objects in **Additional File 5** (BEI), **Additional File 6** (CAMI), and **Additional File 7** (sino-nasal)[82].

The overall diversity of the sinonasal microbiomes collected here were relatively low. Across all specimens, clustering resulted in a total of 300 OTU (plus 6 centroids that were removed before classification by the CHIM2 filter), and the corresponding centroid OTU sequences were classified to 271 named species comprising 150 genera. Although 300 OTU were detected overall, the top 20 OTU comprised 96.7% of reads (**Figure 7**), and only 61 OTU had >50 read counts summed across all >460K primer-match CCS reads. As previously seen, the dominant taxa in the sinonasal microbiome were *Staphylococcus* (OTU_2; see below) and *Propionibacterium acnes* (OTU_1), which together comprised 65.2% of all read counts [35]. Three of the top 20 OTU (and 7 in total) were classified as *Anaerococcus octavius*, which suggests high variation among 16S rRNA genes within this species (**S17 Figure, Additional File 3**).

**Figure 7.** Composition of the sinonasal community. Multiple dots indicate that more than one OTU was classified as the same species. **A**. Overall relative abundance of the top 20 most abundant species. **B**. Number of species observed in 10 or more samples.

We next investigated the relationship between species-level confidence values for each centroid and how many species are shared in the same dbOTU (**S18 Figure, Additional File 3**). This analysis identified several OTU whose centroid assignment belonged to a dbOTU with only 1 or 2 species. These may represent other novel or poorly described species, or alternatively some may represent problems with the taxonomy assignments in the NCBI database.

MED analysis of filtered CCS reads that had been assigned to the high abundance staphylococcal OTU (whose centroid was assigned to *S. epidermidis*) further distinguished among distinct staphylococcal species within the human sinonasal samples, and this was improved when using FL16S compared to V3-V5 reads (**Figure 8**). The presence within the sinonasal communities of additional close relatives to *S. aureus* and *S. epidermidis* clarified how V3-V5 truncated reads likely made some erroneous assignments, compared to FL16S. For example, examination of the trees in **Figure 8** suggests that the V3-V5 ASVs for *S. capitus* and *S. cohnii* are likely misclassified *S. epidermidis* sequences, and also that the *S. hominis* ASV detected with FL16S reads was likely misclassified as *S. lugdunensis* with the V3-V5 reads.

**Figure 8.** Phylogenetic trees of ASVs (MED node representatives) from the human sinonasal community belonging to the *Staphylococcus* OTU, along with staphylococcus NCBI database entries. **A.** FL16S reads and **B.** V3-V5 truncated reads, as in **Figure 4**. Only species detected in one or both dataset are given a non-grey tip color.

### Variation in sinonasal microbial communities among subjects and anatomical sites

Considerable variation in microbial composition was seen among sinonasal specimens, ranging from 3 to 56 OTU per sample and from 50 to 108 per subject. The “core” sinonasal microbiome consisted of 11 OTU that were present across all 12 subjects, whereas most taxa were found in only a few individuals (**Figure 7B**). For subsequent analyses of microbial diversity, absolute counts were normalized to relative abundances after first removing low yield samples and rare taxa, though results were qualitatively similar even with no filtering. We set a minimum sample size of 500 read counts, reducing the number of samples from 122 to 108 (the number of reads collected per sample was highly variable, ranging from 0 to 17,548, mean 3,842 ± 3,229). We also set a minimum OTU size of 50 read counts across the whole set of samples, reducing the number of taxa to only 59 OTU across the dataset. Though rare taxa may play important roles in the sinonasal microbiome, as has been shown in other environments [6], in the absence of dense longitudinal sampling, we could not tell whether these were resident to the sinonasal passages, transients, or contaminants.

Overall, the taxonomic profiles across samples were distinctly more similar within-subjects than within-site, as illustrated by hierarchical clustering and NMDS ordination of samples (**Figure 9, Figure 10A**). This suggests that though the bacterial composition varies at distinct sub-anatomical sites, differences in microbial composition among individuals is much higher.

**Figure 9.** Heatmap of human sinonasal microbiome from 12 subjects. Columns are subjects; rows are species. OTU counts were summed by species-level centroid classification, samples with <500 reads were excluded, then species with <0.2% relative abundance in all samples were dropped. Remaining OTU counts were converted to relative abundances and then log-transformed after adding a pseudocount (1 / # of reads in sample) before hierarchical clustering, showing strong clustering by subject (horizontal colored strip, with different colors indicating the sample’s subject)

**Figure 10.** Diversity of the human sinonasal microbiome by patient, site, and type. **A**. NMDS ordination of log-transformed Euclidean distance matrix of relative OTU abundances in human sinonasal specimens. Some clustering is observed by patient (color), little to no clustering by site (size) or type (shape). **B.** Box-plot of the variation in diversity among sites. The x-axis has all the sites used in the sinonasal community sequencing and the y-axis represents the diversity. Coloring is based on the sample type, swab or biopsy. **C**. OTU richness and Shannon’s effective number of OTU. Box-plot of number of OTUs observed in each patient. The colors are based sample type, swab or biopsy. **D**. Box-plot of Shannon’s effective number of species observed in each patient. The colors are based sample type, swab or biopsy.

Because many OTU were only found in a subset of specimens, we next examined differences in the overall diversity of the samples with respect to subject, anatomical site, and whether obtained by swab or biopsy. Instead of using OTU richness (*i.e.* the total number of OTUs in each sample), we calculated Shannon’s diversity index (which accounts for the relative abundance of distinct OTUs). Analysis-of-variance (ANOVA) of Shannon’s diversity found that, by far, the most important factor accounting for variation in Shannon’s diversity was the subject the sample had come from (**Figure 10B,** p < 2e-16). Sample type (swab versus biopsy) showed no significant effect (p=0.116).

Furthermore, although anatomical site was a significant contributor to the variance, no obvious trends were seen; variation among subjects was much higher (**Figure 10C**, p=0.0082). Swab/biopsy pairs from the same site and subject were extremely similar (p=0.96), indicating no major shift in bacterial composition between the mucosal layer and the tissue immediately beneath, though the latter likely had fewer bacteria overall. These findings were robust to changing the filters used, to using genus- or species-level classifications, and to reformulating the ANOVA model with different factor orders and interaction terms. Furthermore, these results were not an artifact of undersampling in some samples, since there was no correlation between within-sample Shannon’s diversity and sample read count (**S19 Figure, Additional File 3)**. A distinct test that accounts for undersampled rare taxa may be more appropriate (via the breakaway package for R [83]), but due to the relatively low diversity of individual samples, we were unable to apply this test due to a requirement of seeing 6 consecutive frequency classes was not met in any sample). Overall, these results suggest some underlying community structure in the sinonasal cavity, though much of this effect is hidden by the much larger differences in overall microbial composition among subjects.

Finally, to examine whether bacteria might partition differently within the sinonasal cavity, we performed ANOVA on transformed relative abundance measurements for each OTU. Only a single OTU showed a significant effect by anatomical site (*Propiniobacterium acnes*, p=0.009 after Benjamini-Hochsberg FDR correction), and none showed variation by swab versus biopsy. Interestingly, *P. acnes* was least abundant relative to other bacteria in the nasal vestibule (site AB) the largest and most aerated part of the sinonasal cavity, whereas its abundance often made up a major component of the bacterial signature at other less accessible sites (**Figure 10D**).

## Discussion

We report a novel high specificity pan-bacterial molecular diagnostic pipeline for profiling the bacterial composition of microbiome samples, applying amplification and sequencing of full-length 16S rRNA genes (FL16S) with the Pacific Biosciences (PacBio) platform. We exploit circular consensus sequencing (CCS), in which we obtain >10 passes on average of each single molecule sequenced, resulting in CCS reads with exceptionally high quality. This single molecule correction system is not possible on other modern DNA sequencers [19, 84]. Notably, our MCSMRT software is modular and can easily provide inputs to well-established and commonly used downstream microbiome analysis pipelines (namely QIIME and Mothur) at several points before or after OTU clustering, ASV detection, taxonomic assignment, and abundance calculations.

Previous applications of PacBio to sequencing the 16S rRNA gene were initially hampered by higher error rates and insufficient polymerase processivity to leverage circular consensus sequencing [19, 20]. Subsequent improvements in PacBio sequencing chemistry have mostly overcome this [21, 22], and more recent efforts have shown the value of FL16S sequencing by PacBio for and identified the major considerations needed for handling PacBio instead of Illumina 16S reads [23-25]. This work extends and improves upon previous efforts in several ways:

(1) We provide the flexible MCSMRT pipeline to handle processing, clustering, and taxonomic assignment of PacBio 16S reads after first identifying and implementing a series of stringent filters that eliminate many sources of sequencing artifacts, in particular we show that using only the highest fidelity consensus reads for OTU clustering (those with a cumulative expected error, EE, of ≤ 1) effectively eliminates over-calling the number of OTU, which has been a pervasive problem in methods using shorter partial 16S sequences [56], comparable to recent observations using PacBio sequencing of the same BEI mock community as we use here [24].

(2) We generated new PacBio FL16S datasets for pipeline development and benchmarking, including monoculture controls from two species, low and high complexity mock communities, and hundreds of samples from the human sinonasal microbiome.

(3) Because taxonomic assignments remain especially important in the study of human-associated bacteria, we developed a species-level taxonomic classifier for FL16S. To assign taxonomy and confidence values CCS reads, we created a custom-built database constructed from all available FL16S sequences at NCBI, since many commonly used 16S rRNA gene databases lack species-level classifications or lack FL16S genes for many taxa. This allowed us to use a uniform Linneaen hierarchy build a classifier that defined FL16S gene sequences to the species level along with associated confidence values. Our analysis showed improved accuracy and higher confidence when using FL16S sequences compared to partial sequences, which are typical when using short-read Illumina MiSeq 16S survey methods that normally capture only up to ∼500 nt using paired-end sequencing (*e.g.*[63]).

(4) We investigated the use of minimum entropy decomposition (MED) to detect amplicon sequence variants (ASVs) as a way of distinguishing among closely related organisms [45, 85]. This found that decomposing OTUs into ASVs improved identification of closely-related species, although the number of ASVs detected exceeded that expected within the BEI mock community. Some of this could be attributable to intragenomic variation among 16S rRNA gene copies, as seen with our *E. coli* monoculture positive controls, but we also suspect inflated ASV counts due to the particular error profile of CCS reads, which is still biased towards short indel variants, as well as the aggregated effect of true indel variation over the full length of the 16S rRNA gene across bacterial diversity. Future work to improve ASV detection from FL16S CCS reads via MED and/or DADA2 remains ongoing.

We tested our experimental and bioinformatics pipeline on two distinct “mock communities”, showing that use of high-quality CCS reads from FL16S genes has exceptional precision and accuracy at identifying and quantifying the bacteria in complex mixtures. Most impressively, we correctly identified most species in the more complex CAMI community, making only 3 false identifications represented by only 5 CCS reads. Remarkably, we also accurately measured the relative abundance of most species in this complex CAMI community, indicating that our pipeline not only has high taxonomic specificity but high accuracy for quantitative measures of species abundance in complex microbial communities. Underlining the accuracy and precision of our experimental and bioinformatics procedures, 99.81% of filtered reads from monoclonal positive control samples were correctly classified to the species level, and 7 independent negative controls yielded almost no reads that passed our filters.

Following validation of our pipeline using complex mock communities, we applied MCSMRT to the human sinonasal microbiome, finding that the community has a relatively low complexity (with 61 OTU at a frequency of >0.1% across samples); *Staphylococcus* species and *Propionibacterium acnes* dominated across subjects and anatomical sites [35]. Nevertheless, although microbial composition varied much more substantially among subjects than among anatomical sites in the same subject, we nevertheless observed trends in the overall diversity of different sites, with the easily accessible swabs just deep to the nasal vestibule overall reflecting the majority of the healthy sinonasal cavity with the least diverse and least dominated by *P. acnes*. Importantly, we find that swab and biopsy sampling at the same site in the same subject have highly correlated microbial composition, indicating that invasive biopsy sampling is not needed. The large differences among the sinonasal microbiomes among healthy subjects will be of interest in future studies that examine links between sinonasal disease states (particularly chronic rhinosinusitis), bacterial composition, and the innate immune response [86-93]. Our results also show improved discrimination among closely related *Staphylococcus* species when using FL16S compared to the V3-V5 region alone.

PacBio remains more expensive than Illumina per read, though the price has dropped considerably since the introduction of the Sequel instrument and is expected to drop further when higher yield SMRTcells are released. Thus, we expect that the cost trade-off for higher specificity with PacBio (taxonomic and phylogenetic resolution) over higher sensitivity with Illumina (high yields) will rapidly decrease. Our use of primers targeting all nine variable regions maximized specificity, but it may also have narrowed the overall breadth of bacterial diversity we could capture [94, 95], so future studies will investigate primer combinations that maximize both breadth and specificity.

Our results and others show that increasing the length of marker gene sequencing improves the taxonomic and phylogenetic resolution, and we expect that further improvements to sequence processing and analysis will greatly enhance methods that use ASV detection. We further expect that CCS analysis can expand the scope of other marker-based taxonomic and phylogenetic classification schemes, for example, through joint single-molecule sequencing of eukaryotic ITS and 18S rRNA genes from fungi to enrich and extend marker-based databases [96]. Beyond marker gene surveys, metagenomic shotgun sequencing efforts have shown the massive potential for simultaneous profiling of functional gene content and high resolution phylogenetic and taxonomic binning. Although these approaches often remain prohibitively expensive for profiling many host-associated microbiota and may be less amenable to use in clinical diagnostics, we note that metagenomics shotgun assembly and downstream analysis could potentially be greatly enhanced by use of high-quality CCS reads.

## Methods

### Data Availability

MCSMRT https://github.com/jpearl01/mcsmrt,

16S database https://drive.google.com/file/d/1UaWvDnVfGOOtL3ld4BOtl5v7H5igB0To/view

All Sequencing Data as Biosamples (see **Additional File 2, Table S4** and **S8**) https://www.ncbi.nlm.nih.gov/sra/

OTU Tables, Sample Info, and Trees in Phyloseq objects, see **Additional File 5, 6, 7**

#### Ethics statement

Patients were recruited from the Division of Rhinology of the Department of Otorhinolaryngology - Head and Neck Surgery at the University of Pennsylvania with full approval of the Institutional Review Board (Protocol 800614). Informed consent was obtained during the pre-operative clinic visit or in the pre-operative waiting room. Selection criteria for recruitment were patients undergoing sinonasal surgery for non-rhinologic disease entities, *e.g*. pituitary pathology or other cranial base pathologies.

#### Sinonasal sample collection

Sinonasal samples were obtained from patients undergoing sinonasal surgery for non-inflammatory and non-infectious indications (predominately pituitary tumors or other skull base neoplastic process) who had not received antibiotics in the preceding 8 weeks. The Institutional Review Board at The University of Pennsylvania School of Medicine provided full study approval and informed consent was obtained pre-operatively from all patients. Sinonasal specimens were collected as both swabs (S) (BD ESwab collection and transport system) and Tissue (T) (placed in MP lysing matrix tubes). Multiple locations in the sinonasal cavity were sampled including the nasal vestibule (S), inferior turbinate head (T), uncinate process (T), middle meatus (S), maxillary sinus (S)(T), ethmoid sinus (S)(T), superior meatus (S), superior turbinate (T), and sphenoid sinus (S)(T) for a maximum of 12 specimens per patient.

#### DNA extractions

Total DNA was isolated from all samples (swabs and biopsies) using DNeasy Blood & Tissue Kit (Qiagen) according to the manufacturers recommendations with slight modifications. Biopsy material was incubated overnight at 56°C with 570 μl ATL lysis buffer with 30 μl Proteinase K in a Lysing Matrix E tube (MP Biomedicals LLC), homogenized by SPEX 1600 MiniG (Fisher Sci.) for 10min. at 1500 Hz, and centrifuged 1min x 13000 rpm. Swab tubes were treated similarly but initially vortexed for 1min. and spun for 10 seconds and incubated for only 5 min. at 56 °C prior to homogenization. DNA was eluted with 200 μl of the Elution Buffer. DNA quality, and quantity were analyzed by agarose gel electrophoresis and Nanodrop 2000 spectrophotometry.

#### Control DNA samples

The BEI mock community was obtained through BEI Resources, NIAID, NIH as part of the Human Microbiome Project (www.beiresources.org): We used genomic DNA from Microbial Mock Community B (Even, Low Concentration), v5.1L, for 16S RNA Gene Sequencing, HM-782D. The complex CAMI mock community was obtained from the JGI, which had been constructed for the CAMI (Critical Assessment of Metagenomic Interpretation) Hosts Community Challenge for Assessing Metagenomes. Human DNA was isolated from the U937 lymphoblast lung cell line as an off-target control template.

#### FL16S rDNA PCR reactions

Amplifications were performed using 1μl total DNA as template, universal 16S primers F27 and R1492 with four sets of asymmetric barcodes at 0.25 μM (Table S3) [97, 98], and GoTaq Hot Start Master Mix (Promega) or AccuPrime Taq High Fidelity Polymerase with 1μl of 10mM dNTP Mix (Fisher Sci.) in 50μl final volume. Cycling conditions were: 94°C, 3 min; then 22 or 35 cycles of 94°C 30 sec, 54°C 30 sec, 72°C 2 min; following by 5 min final elongation at 72°C. PCR products were cleaned with AxyPrep™ MagPCR (Corning Life Sciences) according to the manufacturer’s protocol and eluted in 40μl of water. Cleaned PCR products were quantified using both using Quant-iT™ dsDNA Assay Kit, high sensitivity (Invitrogen) on BioTek™ FLx800™ Microplate Fluorescence Reader, and AccuClear Ultra High Sesitivity sDNA Quantitation Kit (Biotium). Based on the results, amplicons were normalized to the same concentration prior to pooling amplicons with distinct barcodes into multiplexed sets of 2-4 samples per pool.

#### Pacific Biosciences circular consensus sequencing

Library construction used Pacific Biosciences (PacBio) SMRTbell™ Template Prep Kit V1 on normalized pooled PCR products, and sequencing was performed using the PacBio RS II platform using protocol “Procedure & Checklist - 2 kb Template Preparation and Sequencing” (part number 001-143-835-06). DNA Polymerase Binding Kit P6 V2 was used for sequencing primer annealing and polymerase binding. SMRTbell libraries were loaded on SMRTcells V3 at final concentration 0.0125 nM using the MagBead kit. DNA Sequencing Reagent V4 was used for sequencing on the PacBio RS II instrument, which included MagBead loading and stage start. Movie times were 3 hours for all SMRTcells. PacBio sequencing runs were set up using RS Remote PacBio software and monitored using RS Dashboard software. Sequencing performance and basic statistics were collected using SMRT^®^ Analysis Server v2.3.0.

#### Pre-Clustering Pipeline

MCSMRT accepts CCS data from the PacBio RSII sequencer, and is divided into pre-clustering and clustering steps (**Figure 1, Pre-clustering Pipeline, Additional File 1**). Sequences were generated using the reads of insert (ROI) protocol within Pacific Biosciences SMRT® Analysis Server, reads which had 4 or fewer CCS passes were removed. To further filter low quality or off target sequences, reads failing 3 filters were removed: (a) CCS reads outside the range of 500 to 2000 bp, (b) those that aligned to the hg19 human genome with bwa v0.7.10-r789, and (c) those that did not match both primer sequences with usearch v8.1.1861 [15, 53]. Primer sequences were then trimmed, and reads were oriented 5’ to 3’ with respect to 16S rRNA transcription. Venn diagrams defining read filtration subsets were created using Venny [99].

#### OTU clustering, taxonomic classification of centroids, and OTU abundances

OTUs were generated using the uparse [41] algorithm in the usearch software, using parameters tuned for full-length sequence. In short, reads were de-replicated (and the number, or size, of identical sequences tracked in the header), then sorted by abundance. OTUs were iteratively created at a threshold of 3% divergence from any other existing OTU centroid sequence (*i.e*. reads within 97% similarity to an existing OTU centroid became a member of an existing cluster; otherwise a new OTU was formed with that sequence).

To obtain a database capable of providing a species-level classification of the full-length sequences, all sequences annotated as FL16S genes were downloaded from NCBI in Oct. 2015, and taxonomies were inferred from each read’s 16S gid identifier via the associated txid. This newly formatted database contained species-level taxonomic information for OTU classification (**16S rRNA Microbial Database, Additional File 1**). Representative OTU sequences were assigned a taxonomy using a utax classifier built from this database.

Chimeric sequences were removed during the clustering process based on previously seen OTU centroid sequences (CHIM1 filtering), followed by removal of chimeric centroid OTU using uchime to filter the final OTU sequences using the RDP ‘gold’ sequences [54].

OTU abundance was determined using usearch for filtered reads prior to the expected error threshold, reported as CCS read counts assigned to each centroid OTU. 16S rRNA copy number for the BEI community was estimated from provided quality control data, and OTU abundance by 16S rRNA copy number was calculated in R (Table S1, Figure 8). Expected OTU abundances for the CAMI datasets used the rrndb database to obtain a predicted 16S rRNA gene copy number for each taxon, using the mean of values at the lowest matching taxonomic level.

#### Sub-OTU ordination and phylogenetic methods

A 3% divergence cutoff for OTU clustering is commonly used in comparing various partial 16S fragments [6, 10, 100, 101]. To further examine how individual reads were related to one another, mafft [102] alignments of individual genus/species sequences (including sequences from both OTU and matching NCBI database) were created. Pairwise distance matrices from the alignments were created using ape v4.1 [103] and seqinr v 3.3-6 [104]. 2D non-metric multidimensional scaling (NMDS) ordinations and neighbor-joining trees used vegan v2.4-0 and ggtree v1.8.1 respectively. Data were visualized using ggplot2 v2.2.1 [105-107]. Additionally, maximum likelihood trees (FastTree v2.1.8) were calculated and visualized with ggtree [108].

#### Ecological analyses of the healthy sinonasal microbiome

Measures of ecological diversity (number of species observed, Shannon’s diversity index) were calculated for each sample using vegan before and after filtering to eliminate samples with <500 CCS reads, and OTU with <50 CCS reads across all samples. Count tables were transformed to relative abundances prior to calculating dissimilarity and distance matrices by either Bray-Curtis or Euclidean distance metrics. NMDS ordinations of samples were generated with vegan, and heatmaps created using the gplots package for R.

#### Other Data

Additional miseq data for the same BEI mock community was acquired from [62]. Overlapping paired end reads were joined with COPE, changing default parameters to allow for longer overlap (up to 250 base pairs) [109]. Reads were then imported into MCSMRT and run through default pipeline.

## List of abbreviations

BEI: Biological and Emerging Infections Resources Program
CCS: Circular consensus sequence
CAMI: Critical Assessment of Metagenome Interpretation
EE: Expected Error
JGI: Joint Genome Institute
MCSMRT: Microbiome Classifier using Single Molecule Real-time Sequencing
NCBI: National Center for Biotechnology Information
NIAID: National Institute of Allergy and Infectious Diseases
NMDS: Non-metric Multidimensional Scaling
nt: nucleotide
OTU: Operational Taxonomic Unit
Pacbio: Pacific Biosciences
ROI: Reads of Insert
RDP: Ribosomal Database Project
rrnDB: Ribosomal RNA Operon Copy Number Database

## Declarations

- Ethics approval and consent to participate The Institutional Review Board at The University of Pennsylvania School of Medicine provided full study approval and informed consent was obtained pre-operatively from all patients.
- Consent for publication Not applicable.
- Availability of data and material
- The dataset(s) supporting the conclusions of this article is(are) available in the NCBI SRA repository, [unique persistent identifier and hyperlink to dataset(s) in http://format].
- Software
  - Project name: MCSMRT
  - Project home page: http://github.com/jpearl01/mcsmrt
  - Archived version: DOI or unique identifier of archived software or code in repository (e.g. enodo)
  - Operating system(s): linux 64 bit
  - Programming language: ruby
  - Other requirements: usearch v.8.1.1861
  - License: e.g. GNU GPL, FreeBSD etc.
  - Any restrictions to use by non-academics: none
- For databases, this section should state the web/ftp address at which the database is available and any restrictions to its use by non-academics.

- Competing interests The authors declare that they have no competing interests.
- Funding Funding for this work was provided by R01DC013588 to NAC, R01DC02148, U01DK082316, and NCI HHSN261201600383P to GDE.
- Authors’ contributions
  - Conceived and designed study: JPE, NAC, GDE, JCM
  - Carried out experiments: All
  - Carried out analyses: All
  - Coded software: JPE, AB, RLE
  - Provided materials and reagents: NDA, JNP, NAC, GDE
  - Wrote the manuscript: JPE, JCM, AB, GDE, NAC, JK, SB, RLE, JH
  - Read and approved the final manuscript: All authors

## Acknowledgements

We thank Danielle Reed for useful motivating discussions. We also thank Jason Limbo for technical support, Katherine Kercher for purified genomic DNA from U937 cell line lymphoblast lung from human cells, and Carol Hope for help with preparation of the manuscript. We also thank to Robert Edgar for responsive help with the uparse pipeline. The BEI Bacterial Mock Community was obtained through BEI Resources, NIAID, NIH as part of the Human Microbiome Project Very special thanks to Julio Corral Jr. and Axel Visel at JGI Production Facility for providing the CAMI mock community.

